# TUnA: An uncertainty aware transformer model for sequence-based protein-protein interaction prediction

**DOI:** 10.1101/2024.02.19.581072

**Authors:** Young Su Ko, Jonathan Parkinson, Cong Liu, Wei Wang

## Abstract

Protein-protein interactions (PPIs) are important for many biological processes, but predicting them from sequence data remains challenging. Existing deep learning models often cannot generalize to proteins not present in the training set, and do not provide uncertainty estimates for their predictions. To address these limitations, we present TUnA, a Transformer-based uncertainty aware model for PPI prediction. TUnA uses ESM-2 embeddings with Transformer encoders and incorporates a Spectral-normalized Neural Gaussian Process. TUnA achieves state-of-the-art performance and, importantly, evaluates uncertainty for unseen sequences. We demonstrate that TUnA’s uncertainty estimates can effectively identify the most reliable predictions, significantly reducing false positives. This capability is crucial in bridging the gap between computational predictions and experimental validation.

## Introduction

Characterizing protein-protein interactions (PPIs) is fundamental to understanding many biological processes such as signal transduction, cellular metabolism, and the maintenance of cellular systems.^1^ High-throughput techniques, such as yeast-two-hybrid^2^ and tandem affinity purification^3^, have greatly accelerated identification of PPIs, but these experiments are often time consuming and labor intensive. Recently, deep learning (DL) methods have emerged as a promising alternative.^4^ While protein structure is critical for protein binding, DL models primarily relying on protein sequence, given its relative abundance over structural data, have achieved impressive performance.^5,6^ For example, PIPR^5^ utilizes a siamese recurrent convolutional neural network to capture local and sequential features such as co-occurrence similarity of amino acids and electrostaticity and hydrophobicity based features. PIPR is outperformed in cross-species generalizability by D-SCRIPT,^7^ which combines linear and convolutional layers to learn a predicted contact map for a given protein-protein interaction. Recently, Topsy-Turvy^8^ combined D-SCRIPT with GLIDE,^9^ a network-based approach that considers the local shared-neighbor relationships together with the global network information, and improved cross-species generalizability over D-SCRIPT.

Despite these advancements, a major challenge of DL-based models is its incapability to detect out-of-distribution (OOD) data points and avoid overfitting to training data. Overfitting is especially concerning for PPI prediction, where the vastness of the protein sequence space, and consequently the protein-protein interaction space, cannot be fully captured in the training datasets. Prior state-of-the-art (SoTA) sequence-based models have shown to be effective in predicting PPIs for species they were trained on, but have shown to be poor predictors when tested on untrained species.^5–7^ This limitation in generalizability was further highlighted in a recent study which created a human dataset (referred to as the Bernett dataset) with strategically partitioned training, validation, and test datasets, all with the goal of minimizing sequence similarity and node-degree information.^10^ When evaluated on the Bernett dataset, the PPI prediction models DeepFE, PIPR, D-SCRIPT, and Topsy-Turvy only achieved a balanced accuracy of 0.52, 0.52, 0.50, and 0.56 respectively, underscoring the urgent need for new methods with improved generalizability.^10^

To avoid overfitting to the training data, a powerful strategy is to estimate the uncertainty of the predictions.^11^ Uncertainty awareness in PPI prediction is particularly important because of the huge number of possible protein-protein pairs. Uncertainty aware models provide a measure of confidence alongside its predictions, reporting lower confidence for predictions involving unfamiliar protein pairs or out-of-distribution (OOD) samples, reflecting a self-awareness of its knowledge boundaries.^12^ Uncertainty awareness is particularly important when PPI predictions are used for virtual screening, narrowing down specific protein pairs for experimental validation. Uncertainty estimates can serve as a filter, allowing model users to remove highly uncertain predictions to minimize false positives. As it has yet to be utilized for PPI prediction, the integration of uncertainty awareness into PPI prediction is a novel and necessary advancement, one that could enhance the reliability and applicability of DL in this field.

We have developed **TUnA** (Transformer-based Uncertainty Aware Model for PPI prediction), a sequence-based DL method that leverages implicit structural information for increased generalizability and uncertainty awareness for OOD detection. TUnA has three major components that make up the core framework: ESM-2 protein embeddings, use of Transformer encoder for learning intra and inter protein relationships, and the incorporation of the Spectral normalized neural Gaussian process (SNGP) method for uncertainty awareness.^13^ SNGP enhances deep learning models by applying spectral normalization to the hidden layers and substituting the final fully-connected layer with a Gaussian process (GP) layer. These modifications significantly improve models’ uncertainty awareness while retaining their original predictive accuracy.

We assess TUnA on two widely used benchmark datasets: the cross-species dataset and the Bernett dataset. In the cross-species task, TUnA improves upon the previously best performing Topsy-Turvy as well as the previously benchmarked PIPR^5^ and D-SCRIPT.^7^ Similarly on the Bernett dataset, TUnA is the most accurate and balanced model out of all evaluated methods. Additionally, we show that TUnA’s uncertainty awareness improves calibration and we demonstrate a practical application of uncertainty awareness. Our findings suggest that TUnA not only advances the state of PPI prediction but also is the first model to emphasize uncertainty as a core component.

## Results

### Model overview

The three core components of the TUnA architecture are shown in **Fig. 1**. First, the protein embedding method. Previous works have utilized such as one-hot encoding, hand-crafted physicochemical features, or conjoint triad features.^5,14,15^ Conversely, more recent models such as D-SCRIPT and Topsy-Turvy utilize pre-trained protein language models to embed protein sequences. We utilize the SoTA ESM-2 protein language model which has implicitly learned rich structural information via the masked language modeling objective. While only ever trained on sequence information, ESM-2 can learn structural information as predicting masked residues requires an understanding of evolutionary sequence patterns closely tied to biological structure.^16^ ESM-2 provides a structural information-rich starting point for TUnA.

**Figure 1.**
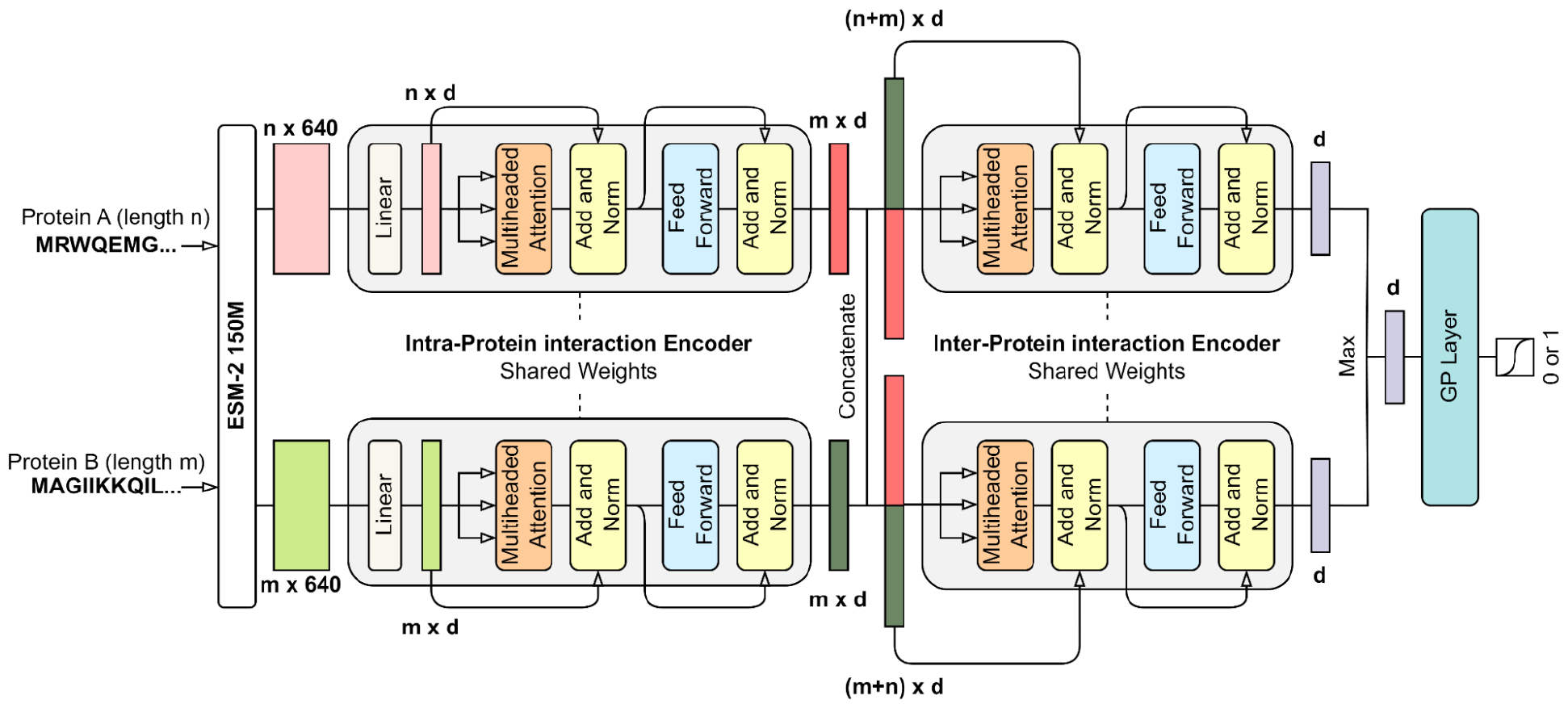
TUnA architecture overview. Protein sequences are embedded using ESM-2 then passed into the intra-protein interaction encoder composed of a linear transformation down to d-dimensions, followed by a standard Transformer encoder. The encoded protein representations are concatenated and passed through the inter-protein interaction encoder. We use both concatenations (A||B and B||A) to ensure permutation invariance. The outputs of the inter-protein interaction encoder are averaged along all non-padded regions to output a d-dimension vector, referred to as the interaction feature vector. The interaction feature vectors are then max-pooled and used as the input for the GP layer. The GP layer returns a uncertainty-adjusted probability used to assign the label the protein pair as interacting or non-interacting.

Second is the Transformer-based architecture. Transformers, widely used in natural language processing for their ability to capture rich long-range dependencies through the multi-headed self-attention mechanism, have shown to be impactful even in drug-protein interaction prediction and protein-protein interaction prediction, as evidenced by TransformerCPI and TransformerGO, respectively.^17–19^

Self-attention can be especially useful for protein sequences as the relationship between residues is not sequential in 3D. For example, a protein may have an important structural motif that is composed of residues distant in the AA sequence but close in the 3D structure. We first use the Transformer encoder twice, once per protein, with the goal of extracting an encoded description of the protein by considering the intra-protein interactions. By concatenating the encoded protein representations and passing it through the inter-protein encoder, the goal is to create an informative representation of the entire protein-protein complex, combining information about the individual proteins and the inter-protein interactions.

Lastly, we implement the Spectral-normalized Neural Gaussian Process (SNGP) method outlined by Liu et al. to introduce distance and uncertainty awareness.^13^ SNGP involves applying spectral normalization to the hidden layers for approximately distance-preserving hidden mappings and replacing the final fully-connected layer with a Gaussian process approximated using random Fourier features. Compared to other uncertainty estimation methods such as Deep Ensembles^20^ and Monte Carlo Dropout,^21^ SNGP only requires only a single network, thus offering a low-cost uncertainty estimate, and also combines the flexibility of neural network models and better uncertainty calibration of GP. During inference, using the learned covariance matrix, TUnA outputs an uncertainty-adjusted probability, p. The uncertainty is a function of p, where uncertainty is highest when p=0.5.

### Performance on cross-species dataset

The cross-species dataset (**Supplementary Table 1**) was constructed by Sledzieski et al., with PPI data originating from the STRING database (v11) filtered to only include experimentally determined physical binding interactions.^7,22^ Furthermore, Sledzieski et al. used CD-HIT^23^ to cluster non-human sequences with human sequences at 40% similarity. Proteins with high similarity to training set proteins were removed to prevent the model from abusing sequence similarity to make predictions.^7^ The datasets are purposely imbalanced, a 10:1 negative to positive ratio, based on the assumption that positive interactions are very rare. Lastly, Sledzieski et al. only include PPIs involving proteins between 50 and 800 AAs.

Following D-SCRIPT and Topsy-Turvy, we report the average and standard deviation of performance metrics across three random initializations. **Fig. 2a** shows the area under precision recall curve (AUPR) and area under receiver operating curve (AUROC) for each model. Given the large class imbalance, AUPR is an appropriate metric for model evaluation. TUnA demonstrates a clear improvement over Topsy-Turvy, achieving the highest AUPR and AUROC scores across all five evaluated species. TUnA achieves higher performance at a significantly reduced computational cost compared to Topsy-Turvy. While Topsy-Turvy required approximately 79 hours for 10 epochs of training, TUnA took approximately 15 hours for 18 epochs. In addition, while Topsy-Turvy utilizes the ProSE^24^ protein language model’s N x 6165 embedding for each length N protein sequence, TUnA uses ESM-2’s N x 640 embedding, requiring about 10 times less memory. In total, the embeddings for the unique mouse sequences for Topsy-Turvy requires ∼367 Gb while TUnA only requires ∼35 Gb. This reduction in computational cost is important for providing a more accessible and practical tool for PPI prediction.

**Figure 2.**
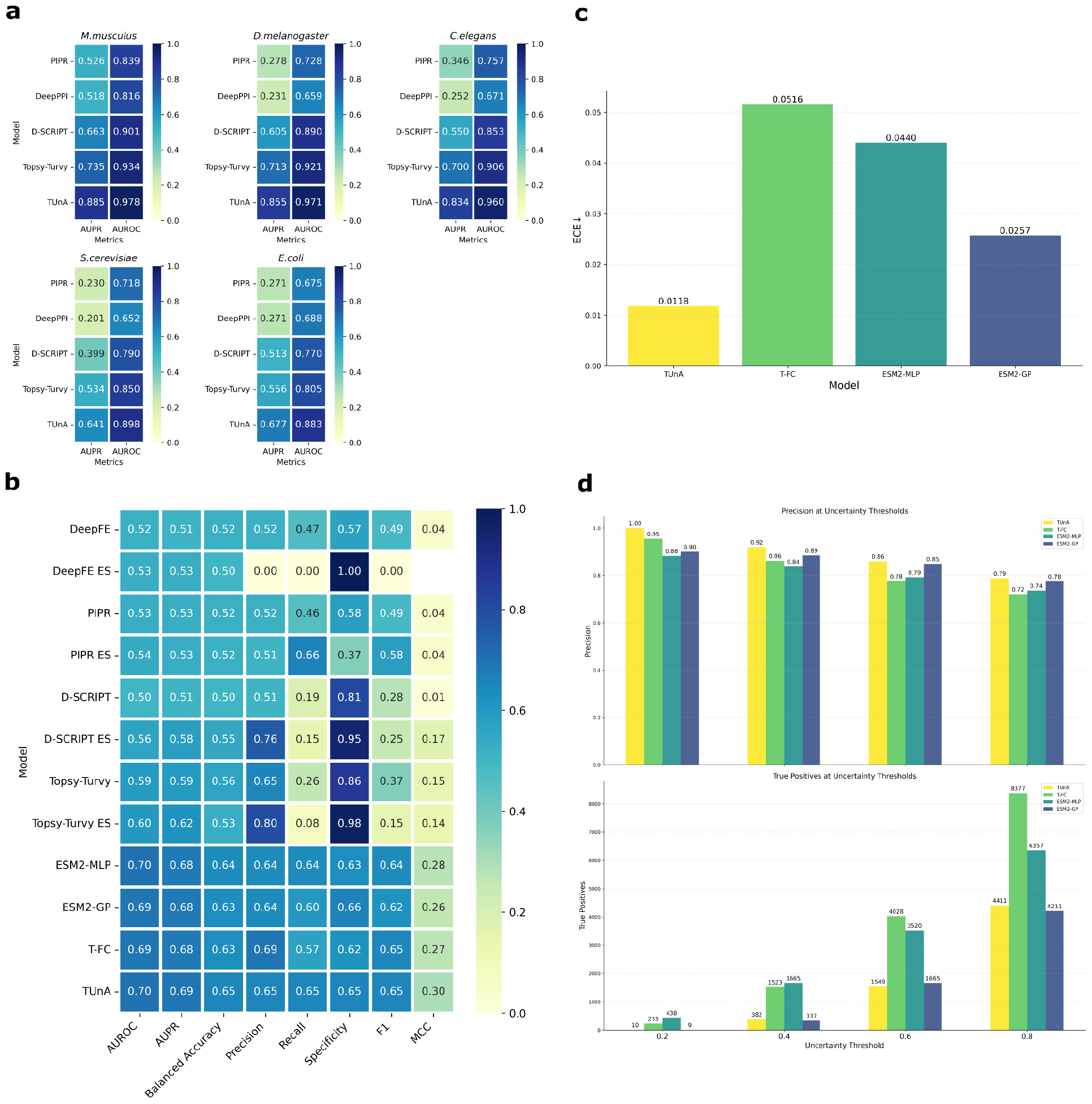
Results on cross-species and Bernett datasets. **a**, Performance on cross-species datasets after being trained on human PPIs. For D-SCRIPT, Topsy-Turvy, and TUnA we report the average over three random initializations. **b**, Performance on the Bernett dataset. ES stands for early stopping, in which model training is stopped when balanced accuracy reaches its peak. Metrics for all models, with the exception of ESM2-MLP, ESM-GP, T-FC, and TUnA, are derived from Bernett et al.^10^ AUROC and AUPR metrics are obtained from the corresponding repository for Bernett et al. We report balanced accuracy, the average of recall and specificity, as was reported by Bernett et al. **c**, Expected calibration error evaluated on the Bernett test dataset post training on the Bernett training dataset. **d**, Precision and number of true positives across different uncertainty thresholds on the Bernett test dataset.

### Performance on Bernett dataset

While the cross-species task can provide a measure of a model’s generalizability, particularly regarding the applicability of knowledge learned from human PPIs to other species, the Bernett dataset^10^ serves as another rigorous benchmark. As a human-only dataset, the Bernett dataset lacks evolutionarily conserved information across species that may inflate model performance. The Bernett dataset aims to minimize data leakage. Starting with positive PPIs from the HIPPIEv2.3^25^ human PPI database, negative PPIs were randomly sampled. Then, the PPIs were partitioned into three parts with the graph partitioning framework KaHIP.^26^ For each partition, an equal amount of negative PPIs was generated by random sampling.

Additionally, CD-HIT was used to reduce redundancy within the partitions, removing any proteins with more than 40% pairwise sequence similarity. Furthermore, to minimize the node degree bias, in which models may predict a PPI based solely on the fact that one of the proteins is frequently involved in positive interactions, the node degree is balanced between the positive and negative examples.

For our experiments, we adhered to the predetermined partitions for training, validation, and testing in the Bernett et al. study.^10^ Due to computational limitations in the protein embedding step, we only used protein sequences with a length of 1500 amino acids or less for the training and validation sets. However, the full test set without this length limitation was used for a fair benchmark comparison. Details of the dataset are shown in **Supplementary Table 2**.

Recently, Sledzieski et al.^27^ showed that ESM-2 embeddings combined with an multi-layer perceptron (MLP) classifier can achieve SoTA performance on the Bernett dataset. Because an official implementation of this model is not publicly available, we train a model closely following Sledzieski et al. ‘s architecture,^27^ referred to as ESM2-MLP, for comparison. To better understand the individual contributions of Transformer encoder and SNGP, we train two additional models. First, we remove the SNGP components from TUnA, removing spectral normalization and replacing the GP layer with a fully-connected layer. We refer to this model as the Transformer with fully-connected (T-FC). Second, we train a model in which we add SNGP to ESM2-MLP, which we refer to as ESM2-GP. For ESM2-GP, we add spectral normalization to ESM2-MLP and replace the last fully connected layer with a GP layer. For T-FC and ESM2-GP, we use the same hyperparameters of their respective original models, except the number of epochs trained for. We determine the number of epochs using the same methodology previously described. The hyperparameters for all models are shown in **Supplementary Table 4, 5**.

We show that TUnA achieves SoTA performance on the Bernett dataset and shows the best balance in the quality and quantity of predictions, evidenced by the highest MCC (**Fig. 2b**). While Topsy-Turvy ES excels in precision and specificity, it has the lowest recall out of all models, suggesting it is heavily biased towards predicting negative interactions. Based on the performances of ESM2-MLP, ESM2-GP, and T-FC we get a deeper insight on the influence of the ESM-2 embeddings, the Transformer encoder, and SNGP.

As previously shown by Sledzieski et al.^27^, ESM2-MLP’s strong performance demonstrates ESM-2 embeddings play a critical role in generalizing to unseen sequences. This aligns with our prior belief that the embeddings may contain implicit learned structural and evolutionary information that are important for predicting PPIs. As unseen sequences are different in sequence but potentially similar in structure, ESM-2 provides additional information leverageable by the model.

Next, a comparison between the no-GP models (ESM2-MLP and T-FC) and GP models (ESM2-GP and TUnA) illustrates the impact of a last GP layer on performance. For the ESM models, a GP layer performs worse performance than MLP. However, for the transformer models, TUnA has better overall performance than T-FC. The difference in response to the GP underscores the GP’s sensitivity to the input features used to update the covariance matrix. Given the Transformer encoder’s capacity to generate more informative interaction feature vectors compared to the MLP, incorporating a GP with T-FC (leading to TUnA), is a more lucrative strategy than incorporating GP to ESM-MLP. We note that the inputs to the GP are the same dimension in both ESM-GP and TUnA, suggesting the difference is not due to a difference in dimensionality but rather the quality of the input features. Thus, we believe the higher computational cost of the Transformer is justified when used in conjunction with SNGP.

### Effect of uncertainty awareness

We highlight our second novel contribution to the field of PPI prediction–uncertainty awareness. In general, DL models can be overconfident for unseen and OOD data.^28^ Overconfidence results in misleading and unreliable predictions. Thus, SNGP is a core part of TUnA, enabling it to make predictions reflecting its knowledge and confidence. Confidence calibration, measured by the expected calibration error (ECE)^29^ assesses the quality of a model’s uncertainty awareness, where better calibration results in a lower ECE. The Bernett test set, given it has no sequences seen during training, is OOD with respect to sequence and thus ideal to evaluate the models’ response to OOD data. We calculate the ECE for best performing models TUnA, T-FC, ESM2-MLP, and ESM2-GP.

While **Fig. 2b** suggests T-FC, ESM2-MLP, and ESM2-GP have comparable performance metrics, their respectives ECEs reveal a drastic difference in calibration. As shown in **Fig. 2c**, models without SNGP (T-FC and ESM2-MLP) have significantly higher ECEs compared to their counterparts with SNGP (TUnA and ESM2-GP). These results suggest SNGP can be an effective method for adding uncertainty awareness to PPI prediction models. Additionally, we observe a similar pattern seen in **Fig. 2b** where the improvement in ECE between T-FC and TUnA is greater than the improvement between ESM2-MLP and ESM2-GP, further justifying and highlighting the advantage of the Transformer-GP combination. Overall, TUnA is the best calibrated and most uncertainty aware model, evidenced by the lowest ECE.

In a study involving protein engineering, Parkinson et al. use uncertainty to select and narrow down the most certain predictions for experimental evaluation to save cost and time.^30^ While we cannot experimentally validate predictions in this study, we describe a practical application of uncertainty awareness using the Bernett test set. For selecting experimental candidates, precision is a key metric considering the number of false positives must be minimized. In other words, the quality of the predictions can be a more important factor than the number of predictions. For TUnA, T-FC, ESM2-MLP, and ESM2-GP, we calculate the precision after removing predictions above the uncertainty thresholds 0.2, 0.4, 0.6, and 0.8, where 0.2 represents the most stringent threshold. For all models, we use the predictive uncertainty, defined as (1−*p*)(*p*)/0.25, where p is the probability of interaction. In addition, we count the number of true positives within each threshold.

In all models, the precision increases as we remove uncertain predictions (**Fig. 2d**). Furthermore, the models incorporating SNGP see a higher precision across different thresholds compared to the models without, TUnA notably having perfect precision at the 0.2 threshold. In reality, downstream validations are often time consuming where the precision of model predictions is the top priority and TUnA is particularly useful for filtering out predictions based on uncertainty.

## Discussion

We introduced TUnA, a novel uncertainty aware sequence-based model for PPI prediction. TUnA utilizes ESM-2 embeddings as well as Transformer encoders for extracting intra- and inter-protein interactions. In addition, we incorporated SNGP to add uncertainty awareness. To the best of our knowledge, TUnA is the first method to incorporate uncertainty awareness to the PPI prediction task.

First, we showed TUnA improves upon the existing methods for predicting cross-species PPIs as well as for predicting human PPIs without sequence similarity-based data leakage, node degree bias, or any evolutionarily conserved information available in the cross-species task. Second, we explored TUnA’s uncertainty awareness as well as the contributions of the ESM-2 embeddings, the Transformer encoder, and SNGP in model performance. We compared TUnA against three different models all utilizing ESM-2 embeddings, T-FC, ESM2-MLP, and ESM2-GP. We found ESM-2 embeddings are a rich starting point for any model given its implicitly learned structural and evolutionary information, even at the 150M parameter level. However, the differences in the models became more apparent when looking at each model’s level of uncertainty awareness, measured by the ECE. Incorporation of SNGP improved uncertainty awareness in all cases but more significantly for the Transformer-based model than the MLP-based model, highlighting the advantage of the Transformer-GP combination. We demonstrated that uncertainty awareness is not only a theoretical advantage but has practical implications for improving precision and reducing the risk of false positives, which is crucial for selecting targets for the follow up experimental validations.

In conclusion, TUnA represents an advancement of the PPI interaction prediction field as a state-of-the-art, uncertainty aware method. Through uncertainty awareness, we hoped to bridge the gap between computational predictions and practical application–TUnA’s uncertainty estimates provide a simple and effective way for anyone to select the most promising PPI candidates. Future work can continue to improve upon robustness against unseen sequences, continuing to push the boundary of what is possible with sequence alone.

## Methods

TUnA is an end-to-end framework designed to process two embedded protein sequences and output a binary prediction indicating non-interaction (0) or interaction (1) as well as a corresponding uncertainty estimate between 0 and 1, where 1 represents the highest possible uncertainty. Following Liu et al.’s methodology^13^, we apply spectral normalization to the model’s weights. Spectral normalization divides the hidden layer weights by their largest singular value, regularizing the amount of stretching or compression carried out by the hidden layers, ensuring approximately distance-preserving hidden mappings crucial for the TUnA’s uncertainty awareness. Additionally, we replace all instances of the ReLU activation function with the Swish activation function, as Ramachandran et al. have shown its effectiveness over ReLU.^31^

### Protein Embedding

We utilize the ESM-2 pre-trained protein language model to transform protein sequences into vector representations. While ESM-2 offers a range of pre-trained models, varying in size from 8M to 15B parameters, we opt for the 150M parameter model due to computational limitations. Given an amino acid (AA) sequence with length N, the embedded representation is an N x 640 matrix. Given that residue distances in sequence are not reflective of the residue distances in 3D structure, positional embeddings are not added.

### Intra-protein feature extraction

To capture the intra-protein interactions, we individually process each protein in the input pair with a Transformer encoder. The protein sequences, represented as N x 640 matrices, are first projected into N x d matrices, where d is the hidden dimension. A mask is applied to the padded regions such that padded regions are ignored during the self-attention block. The output of the intra-protein encoder is a set of encoded N x d representations for each protein, capturing essential intra-protein relationships and features.

### Inter-protein feature extraction

While the original Transformer decoder is tailored for text generation, our PPI prediction task requires feature extraction rather than sequential generation. Therefore, we propose the use of a secondary encoder in place of the decoder. This inter-protein feature encoder takes as input the concatenated encoded representations from the intra-protein feature extraction step. For instance, if Protein A has length N and Protein B length M, their encoded outputs would be N x d and M x d matrices, respectively. Consequently, the input for the inter-protein feature extraction module becomes a (N+M) x d matrix.

The concatenated sequence is treated as one entity, allowing each residue to weigh the attention across residues from both proteins. This reflects the biological nature of protein interactions, which often involve a combination of domains and motifs from both proteins. The inter-protein encoder returns a (N+M) x d matrix as the output. Lastly, we average over the real parts of the sequence (all non-padded regions) resulting in a d dimension vector, which we refer to as the interaction feature vector.

To ensure permutation invariance it is important that the final prediction should not depend on the order of the input proteins (Protein A, Protein B vs Protein B, Protein A). To do so, we process both concatenated sequences (A||B and B||A) through the inter-protein encoder. With the resulting two interaction feature vectors, we apply max pooling such that our final d-dimensional vector, the symmetric interaction feature vector, will be consistent regardless of the protein sequence input order. This permutation invariance is crucial for consistent and predictable PPI predictions.

### Gaussian Process prediction module

As outlined by Liu et al.,^13^ the standard final fully-connected layer is replaced by a Gaussian process layer which is conditioned on the symmetric interaction feature vectors during the final epoch of training. The kernel is approximated by the random Fourier features approximation.^32^ At the last epoch of training, we calculate the covariance matrix, allowing us to generate both a mean logit and its variance for each example during evaluation. Using the mean and variance, the uncertainty-adjusted probability, p, is calculated using the mean-field approximation.^13^ Following Liu et al., we define uncertainty as (1−*p*)(*p*)/0.25, meaning uncertainty is the highest when p=0.5, indicating an OOD sample. To understand why, note that a GP with a radial basis function kernel begins with a prior mean of zero, updating this belief according to the training data.^33^ For out-of-distribution (OOD) samples, the GP reverts to its prior mean of 0, leading to an output logit of zero. Since sigmoid(0) equals 0.5, distant examples are expected to yield a predicted probability *p* of 0.5 (**Supplementary Fig. 1)**. We use the uncertaintyAwareDeepLearn 0.0.5 library (https://github.com/Wang-lab-UCSD/uncertaintyAwareDeepLearn) to implement the last layer GP.

### Implementation and training details

We use the code for TransformerCPI,^18^ a Transformer-based protein-drug interaction prediction model, as a starting point and heavily adapt the architecture and workflow.^18^ TUnA is implemented in PyTorch 1.13.1 with CUDA 11.6 and trained on a NVIDIA A6000 with 48 GB of memory. TUnA minimizes the binary cross-entropy loss with the Adam + Lookahead optimizer.^34,35^ While Adam does not require a learning rate scheduler, we observed adding a StepLR scheduler improved performance. To determine the number of epochs, we trained the TUnA for 20 epochs, then re-trained TUnA until the epoch when it achieved the lowest validation loss to minimize overfitting. Given the large number of hyperparameters and consequently the computational cost of traditional grid search, we identify the most important hyperparameters based on early validation performance. As the size of the hidden dimensions appeared to have a large impact on performance, we identified the optimal hidden dimension through grid search (**Supplementary Table 3**). The selected hyperparameters for TUnA are described in **Supplementary Table 4**.

During training, we limit the maximum sequence length to 512 AAs due to computational limitations. If a sequence exceeds this length, we randomly select a continuous 512 AA-long subset for each training instance, ensuring varied exposure to different sequence regions. Because we train using mini-batches, we zero-pad sequences shorter than 512 AAs. We note that this is only applied during training, and that during testing, the model considers the entire sequence.

## Data availability

All data used in this paper is deposited and publicly available at https://github.com/Wang-lab-UCSD/TUnA

## Code availability

Source code used to generate all results shown in this paper is deposited and publicly available at https://github.com/Wang-lab-UCSD/TUnA

## Acknowledgements

This project is partially supported by NIH (R21AI158114 and R01AI150282).

## Author Information

### Contributions

W.W. proposed the project and supervised along with J.P. Y.K., J.P., and C.L. wrote code for the model.

Y.K. performed model training and evaluation. All authors analyzed the results and wrote the manuscript.

## Ethics declarations

### Competing interests

The authors declare no competing interests.

## Notes

### Competing Interest Statement

The authors have declared no competing interest.

