## Supplemental Information for "TUnA: An uncertainty aware transformer model for sequence-based protein-protein interaction prediction"

##### **Supplementary Information**

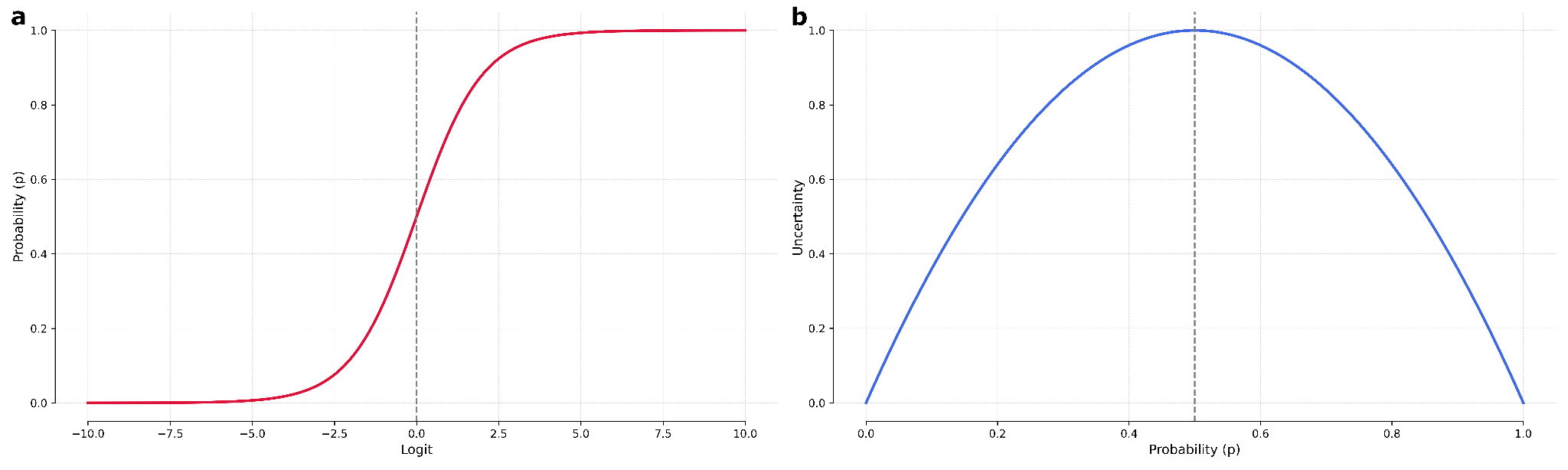

**Supplementary Figure 1. a,** The sigmoid function curve. When the logit is 0, the probability is 0.5. **b,** The uncertainty curve, defined as u(x) = (1−*p*)(*p*)/0.25. When the probability is 0.5, the uncertainty is 1, the highest possible value.

**Supplementary Table 1**. Cross-species dataset breakdown by species.

| Species | # Negative Interactions | # Positive Interactions | # Total Interactions | # Unique Sequences |
| --- | --- | --- | --- | --- |
| Human (Train) | 383,448 | 38,344 | 421,792 | 15,816 |
| Human (Validation) | 47,931 | 4,794 | 52,725 | 15,525 |
| Mouse (Test) | 50,000 | 5,000 | 55,000 | 37,497 |
| Fly (Test) | 50,000 | 5,000 | 55,000 | 19,213 |
| Roundworm (Test) | 50,000 | 5,000 | 55,000 | 25,429 |
| E. coli (Test) | 20,000 | 2,000 | 22,000 | 7,138 |
| Yeast (Test) | 50,000 | 5,000 | 55,000 | 5,664 |

**Supplementary Table 2.** Bernett dataset breakdown.

| Species | # Negative Interactions | # Positive Interactions | # Total Interactions | # Unique Sequences |
| --- | --- | --- | --- | --- |
| Train | 81,596 | 81,596 | 163,192 | 4,286 |
| Train (1500 or less) | 64,781 | 64,811 | 129,592 | 3,894 |
| Validation | 29,630 | 29,630 | 59,260 | 3,711 |
| Validation (1500 or less) | 26,772 | 26,893 | 53,665 | 3,543 |
| Test | 26,024 | 26,024 | 52,048 | 3,022 |

**Supplementary Table 3.** Hidden dimension grid search results for cross-species and Bernett datasets.

| Cross-species |  |  |  |  |  |  |  |  |
| --- | --- | --- | --- | --- | --- | --- | --- | --- |
| Hidden dim. | AUROC | AUPR | Accuracy | Precision | Recall | Specificity | F1 | MCC |
| 64 | 0.982 | 0.916 | 0.973 | 0.886 | 0.806 | 0.990 | 0.844 | 0.830 |
| 128 | 0.984 | 0.922 | 0.975 | 0.883 | 0.834 | 0.989 | 0.857 | 0.844 |
| 256 | **0.986** | **0.932** | **0.976** | **0.886** | **0.848** | **0.989** | **0.867** | **0.854** |
| Bernett |  |  |  |  |  |  |  |  |
| Hidden dim. | AUROC | AUPR | Accuracy | Precision | Recall | Specificity | F1 | MCC |
| 64 | **0.666** | **0.665** | **0.622** | **0.629** | 0.598 | **0.646** | 0.613 | **0.244** |
| 128 | 0.663 | 0.660 | 0.616 | 0.613 | 0.636 | 0.597 | 0.624 | 0.233 |
| 256 | 0.656 | 0.656 | 0.610 | 0.596 | **0.688** | 0.532 | **0.639** | 0.222 |

Performance is evaluated on respective validation datasets, using the epoch with the lowest validation loss, based on a single initialization with random seed 47.

**Supplementary Table 4.** TUnA hyperparameters for the cross-species and Bernett dataset tasks

| Model Hyperparameters | Cross-species dataset | Bernett dataset |
| --- | --- | --- |
| Max. sequence length | 512 | 512 |
| Hidden dimension | 256 | 64 |
| Number of layers | 1 | 1 |
| Number of heads | 8 | 8 |
| Dropout | 0.2 | 0.2 |
| Feedforward dimension (4x hidden dim) | 1024 | 256 |
| Number of GP Random fourier features | 4096 | 4096 |
| Batch size | 16 | 16 |
| Epochs | 18 | 14 |
| Random seed | 47, 94, 141 | 47 |
| Optimizer/scheduler Hyperparameters |  |  |
| Learning rate | 0.0001 | 0.0001 |
| Weight decay | 0.00001 | 0.00001 |
| Lookahead alpha | 0.8 | 0.8 |
| Lookahead k | 5 | 5 |
| StepLR step size | 2 | 2 |
| StepLR gamma | 0.93 | 0.93 |

Note: Feedforward dimension is 4x the hidden dimension, per original Transformer paper.

**Supplementary Table 5.** Hyperparameters for T-FC, ESM2-MLP, and ESM2-GP

| Model Hyperparameters | T-FC | ESM2-MLP | ESM2-GP |
| --- | --- | --- | --- |
| Max. sequence length | 512 | No limit | No limit |
| Hidden dimension | 64 | 64 | 64 |
| Number of layers | 1 intra/inter encoder layer | 2 hidden layers | 2 hidden layers |
| Number of heads | 8 | N/A | N/A |
| Dropout | 0.2 | 0.1 | 0.1 |
| Feedforward dimension | 256 | 64 | 64 |
| Number of GP Random fourier features | 4096 | N/A | N/A |
| Batch size | 16 | 16 | 16 |
| Epochs | 5 | 9 | 10 |
| Random seed | 47 | 47 | 47 |
| Optimizer/scheduler Hyperparameters |  |  |  |
| Learning rate | 0.0001 | 0.0001 | 0.0001 |
| Weight decay | 0.00001 | 0.00001 | 0.00001 |
| Lookahead alpha | 0.8 | 0.8 | 0.8 |
| Lookahead k | 5 | 5 | 5 |
| StepLR step size | 2 | 2 | 2 |
| StepLR gamma | 0.93 | 0.93 | 0.93 |
